# Heterogeneity of the human immune response to malaria infection and vaccination driven by latent cytomegalovirus infection

**DOI:** 10.1101/2024.04.28.591548

**Authors:** Reena Mukhiya, Wim A. Fleischmann, Jessica R. Loughland, Jo-Anne Chan, Fabian de Labastida Rivera, Dean Andrew, James G. Beeson, James S McCarthy, Bridget E Barber, J. Alejandro Lopez, Christian Engwerda, Richard Thomson-Luque, Michelle J Boyle

## Abstract

Human immune responses to infection and vaccination are heterogenous, driven by multiple factors including genetics, environmental exposures and personal infection histories. For malaria caused by *Plasmodium falciparum* parasites, host factors that impact on humoral immunity are poorly understood. We investigated the role of latent cytomegalovirus (CMV) on the host immune response to malaria using blood stage *P. falciparum* Controlled Human Malaria Infection (CHMI) and in a MSP1 vaccine Phase 1a clinical trial. CMV seropositivity was associated with reduced induction of parasite specific antibodies following malaria infection and vaccination. During infection, reduced antibody induction was associated with modifications to the T -follicular helper (Tfh) cell compartment. CMV seropositivity was associated with a skew towards Tfh1 cell subsets before and after malaria infection, and reduced activation of Tfh2 cells. Protective Tfh2 cell activation was only associated with antibody development in CMV seronegative individuals, and a higher proportion of Tfh1 cells was associated with lower antibody development in CMV seropositive individuals. During MSP1 vaccination, reduced antibody induction in CMV seropositive individuals was associated with CD4 T cell expression of terminal differentiation marker CD57. These findings are particularly relevant for malaria endemic regions where CMV infection is acquired early in life and may modify immunity to malaria gained during infection or vaccination.

## Introduction

The human immune response to infection and vaccination is heterogenous. Responses range from robust responses that resolve infection, to sub-optimal responses that may lead to severe disease or fail to generate long-term protection. This heterogeneity reflects the interplay of genetics, environment, and personal exposure histories. For malaria caused by *P. falciparum* parasites, infection in endemic areas ranges from asymptomatic parasitemia to severe disease and death [1]. At the population level, the largest disease burden occurs in children under the age of five, who develop immunity after repeated infection throughout childhood. However, the rate of immune development is heterogeneous, with some children rapidly gaining protection while others experience numerous symptomatic episodes, despite similar exposure [2]. Heterogeneity is also seen in responses to malaria vaccination, both between individuals within malaria endemic areas, and between populations across geographical locations [3–5]. Of note, responses to *P. falciparum* malaria vaccines are consistently lower in low- and middle-income countries (LMIC) compared to those reported during Phase 1 trials in high income countries. For example, antibodies to circumsporozoite protein (CSP) following experimental vaccination with radiation-attenuated whole sporozoite parasite vaccine (PfSPZ) were significantly lower in Tanzanian and Malian adults, compared to adults in the USA [6]. Further, CSP antibodies induced in Kenyan adults with the licenced malaria subunit vaccine RTS,S are generally lower than those seen following vaccination of adults in the USA [7–10]. As such, dissecting factors that modify immune responses to *P. falciparum* may inform development of second-generation vaccines with better protection, or other avenues for malaria control.

Immunity to malaria generated by infection or vaccination is mediated by antibodies that prevent parasite replication, largely via Fc mediated mechanisms [11,12]. Antibody induction is supported by the CD4^+^ T cell compartment, particularly T follicular CD4^+^ T (Tfh) cells, which activate B cells to drive germinal centre activation and antibody development [13], including during malaria [14]. In human infection, Tfh cell subsets can be identified based on CXCR3 and CCR6 chemokine expression, and specific Tfh cell subsets are associated with the development of antibodies following infection and vaccination in a context dependent manner [15]. We have previously evaluated the role of Tfh cells in antibody development during *P. falciparum* malaria, using the powerful controlled human malaria infection (CHMI) model that allows the longitudinal analysis of the immune response to a single infection without the confounding effects of prior malaria exposure and heterogeneity in timing of infection that occur when investigating immunity in naturally exposed individuals [16]. We showed that early activation of Tfh2 cells was associated with induction of functional antibodies, and Tfh1 cells were associated with short lived plasmablasts which have negative roles in germinal centre formation [17,18]. While age has been shown to influence Tfh2 and Tfh1 proportions and activation [19,20], other host factors that influence Tfh cells during malaria are unknown.

One of the multiple factors known to modify the human immune response is cytomegalovirus (CMV) infection, a ubiquitous beta-herpes virus which establishes a life-long persistence and results in major remodelling of the immune response [21]. In monozygotic twins with discordant CMV serostatus, over 50% of immune parameters tested were influenced by CMV seropositivity [22], with impacts across the immune landscape, including monocytes, NK cells, and T cells [23]. Within the T cell compartment, CMV seropositivity has been associated with the expansion of memory cells that lose expression of costimulatory receptors CD27 and CD28, and gain expression of markers of terminal differentiation including CD57 [24]. While these changes are most striking in CD8^+^ T cells, they also occur within CD4^+^ T cells [25]. This immune remodelling has consequences for subsequent pathogenesis of a broad range of diseases and induction of antibody following infection with other pathogens [26]. The global seroprevalence of CMV is 83%, which ranges from less than 50% in high income countries to up to 100% in LMIC [27], where the majority of infants are infected in the first year of life [28]. These geographical differences in CMV seroprevalence have implications for vaccine efficacy for pathogens where CMV can modify the immune response [29]. Whether CMV can modulate the host immune response to malaria infection or vaccination is unknown.

To investigate the impact of CMV serostatus on antibody induction during malaria infection and after vaccination, we investigated antibody induction in CMV seronegative and seropositive individuals enrolled in CHMI studies [17] and in a phase 1a clinical trial of a full-length MSP1 vaccine candidate [30,31]. We found that CMV seropositivity was associated with reduced antibody induction following both CHMI and vaccination, and that these reductions were associated with remodelling of the CD4+ T cell compartment. These findings have implications for the development of malaria immunity, either induced by infection or vaccination, in children in endemic areas where CMV infection occurs in infancy, and CMV seropositivity reaches 100% [28,32].

## Results

### Latent CMV infection is associated with reduced antibody induction following controlled human malaria infection in adults

To investigate the impact of latent CMV infection (quantified as CMV seropositive) on the immune response to *P. falciparum* infection, we analysed antibody and Tfh responses in 40 malaria-naïve adults during blood stage CHMI (median age 25.5, range 18-52 years) [17]. Within the cohort, 21/40 (52%) individuals were seropositive for CMV. Sex, Epstein-Barr virus (EBV) seropositivity and age, were comparable between the CMV negative and positive individuals (Table 1).

**Table 1:**
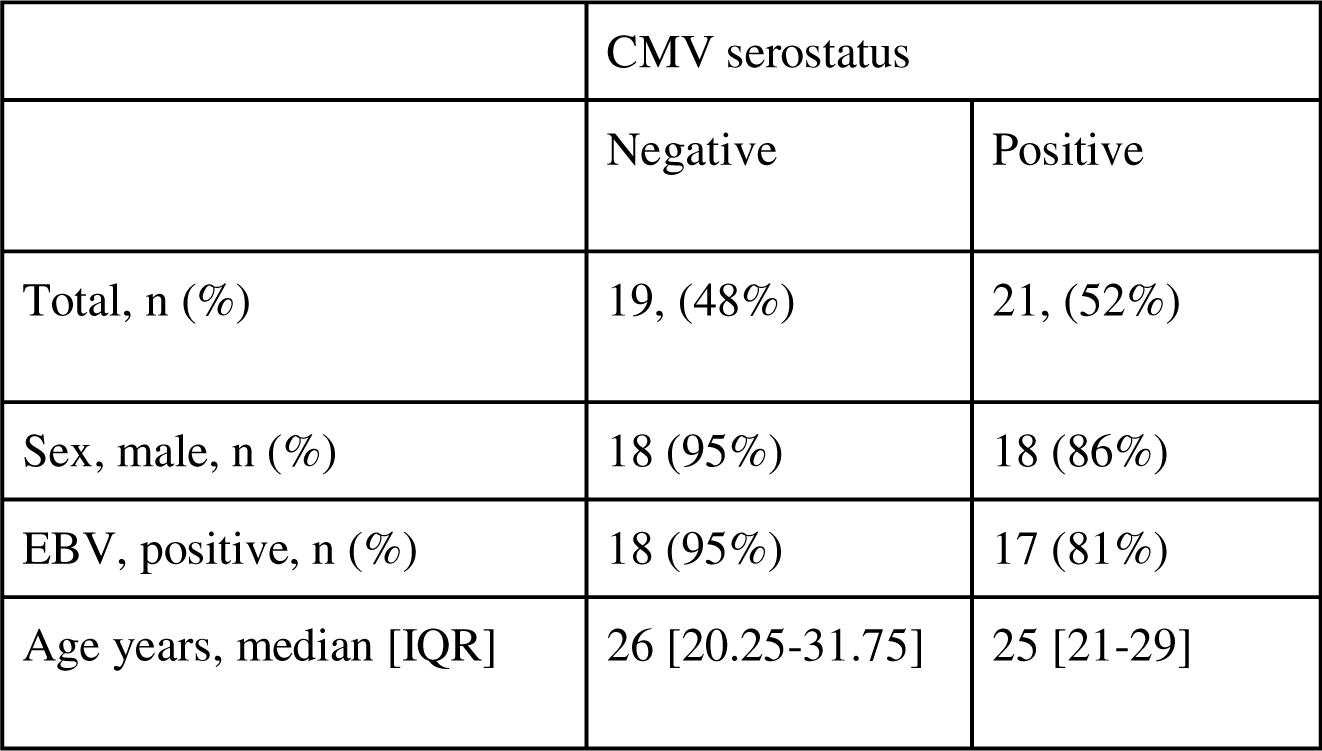
Study populations demographics.

|  | CMV serostatus |  |
| --- | --- | --- |
|  | Negative | Positive |
| Total, n (%) | 19, (48%) | 21, (52%) |
| Sex, male, n (%) | 18 (95%) | 18 (86%) |
| EBV, positive, n (%) | 18 (95%) | 17 (81%) |
| Age years, median [IQR] | 26 [20.25-31.75] | 25 [21-29] |

As previously reported, the induced antibody response was measured as an antibody score, that captured the breadth, magnitude and functionality of antibodies to the merozoite parasite stage, and major merozoite antigen, merozoite surface protein 2 (MSP2) [17]. Antibody score was significantly lower in CMV seropositive individuals (Fig. 1 A). There was no difference in antibody score with EBV status, and antibody score was not associated with age (Fig. 1 B-D).

**Figure 1.**
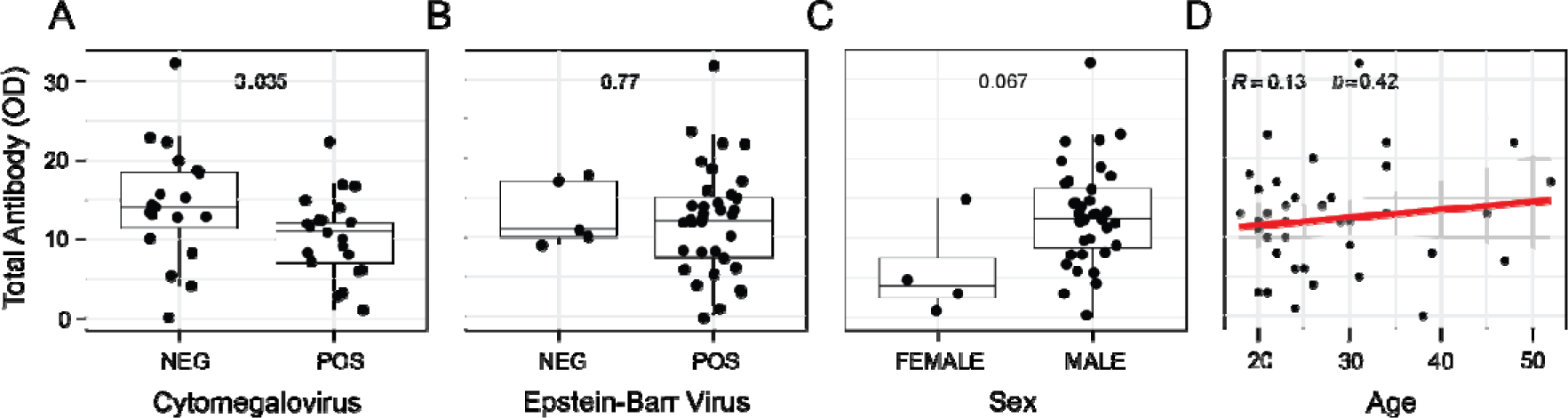
Antibodies induced by controlled human malaria infection are reduced in adults with latent CMV infection. IgM, IgG and functional antibodies to merozoite and major merozoite antigen MSP2 were quantified and antibody score used to capture the total magnitude, breadth and functionality of the responses. Antibody score stratified by **(A)** CMV serostatus (CMV-n=19, CMV+ n=21), **(B)** EBV serostatus (EBV– n=5, EBV+ n=35), **(C)** Sex (female n=3, male n=37), and **(D)** correlated with age. A-C data is Tukey boxplots with the median, 25^th^ and 75^th^ percentiles. The upper and lower hinges extend to the largest and smallest values, respectively but no further then 1.5XIQR from the hinge. P are Mann-Whitney U test. D, rho and P are Spearman’s correlations.

### CMV seropositive adults have reduced induction of antibodies following controlled human malaria infection

To assess the antibodies that were contributing to reduced antibody score in CMV positive individuals, IgM, IgG (total and subclasses) and functional antibodies targeting the merozoite were analysed. In CMV positive individuals, IgG to the merozoite was lower at end of study (EOS, Day 27-36), but there were no differences in the magnitude of induced IgM (Fig. 2 A). The reduced IgG in CMV positive individuals was driven by a reduced IgG1 response. The magnitude of IgG2 and IgG3 were also lower, however differences were not statistically significant (Fig. 2 B). IgG1 is a cytophilic antibody which has important functional capacity to fix complement and interact with Fc receptors on phagocytes. These functional antibody responses have essential roles in immunity to malaria and can target the merozoite to block invasion of the red blood cell and mediate protection [11,33,34]. Consistent with reduced IgG1 induction in CMV positive individuals, CMV positive individuals had reduced induction of antibodies that could fix complement (measured by C1q fixation, the first step in the classical complement cascade [33]), and reduced binding of FcγRII and FcgγIII, which are involved in phagocytosis of parasites by neutrophils and other immune cells [35] (Fig. 2 C/D). However, antibodies that could mediate opsonic phagocytosis by the THP-1 pro-monocytic cell line, which primarily involves FcγRI [35], did not differ between the two groups (Fig. 2 E). There was no difference in the antibody response to MSP2 (Supplementary Fig. 1). Together these data show that CMV is an important modulator of the primary immune response to malaria infection.

**Figure 2.**
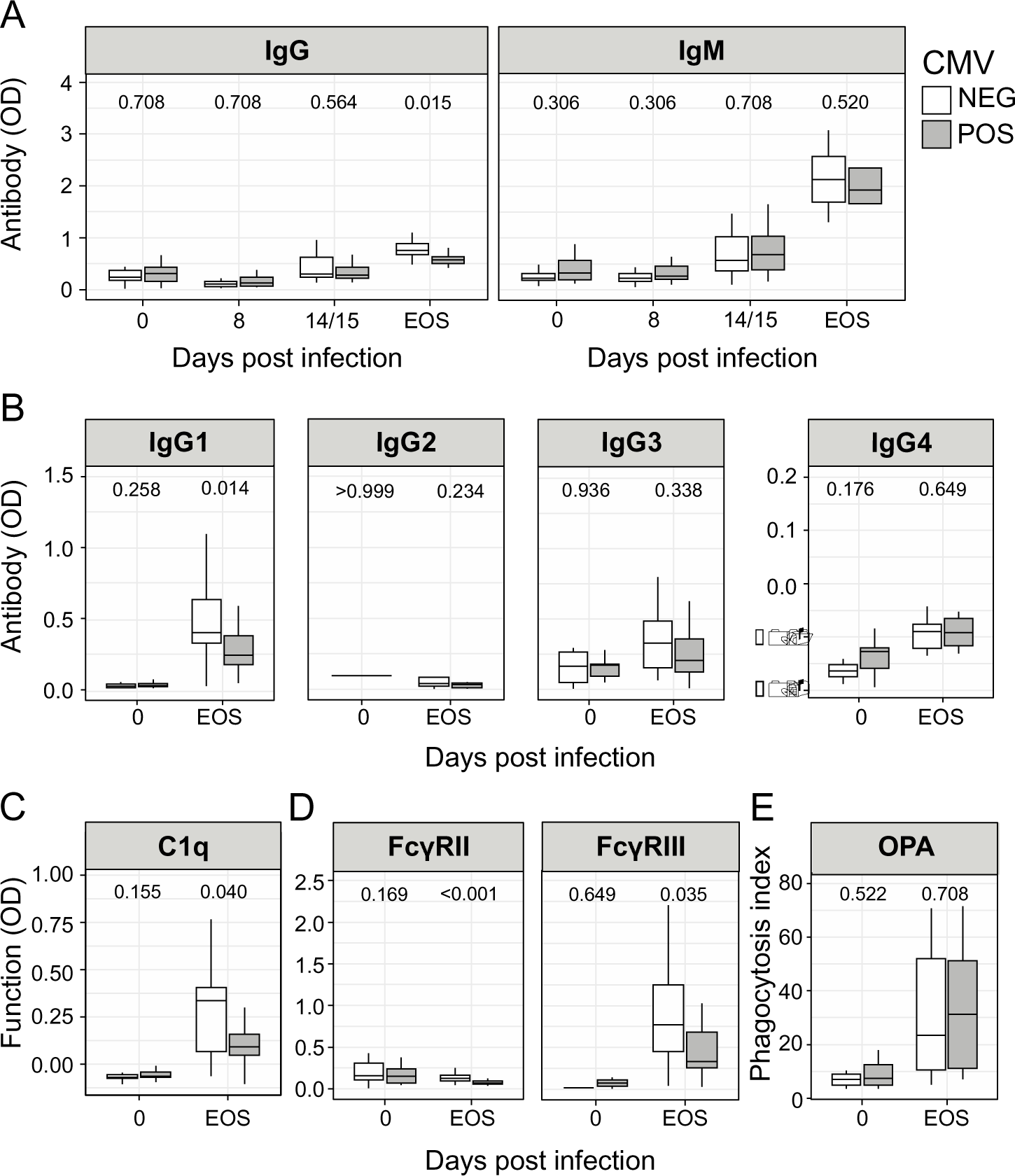
IgG1 and functional antibodies to the merozoite induced by controlled human malaria infection are reduced in CMV infected adults. **(A)** IgG and IgM antibodies, **(B)** IgG subclass antibodies and levels of functional antibodies that can **(C)** fix C1q complement component, **(D)** cross link FcγRII and FcγRIII, and **(E)** drive opsonic phagocytosis by THP1 cells, targeting merozoite stage parasites, stratified by CMV serostatus. CMV-white bars (n=19), CMV+ grey bars (n=21), data is Tukey boxplots with the median, 25^th^ and 75^th^ percentiles. The upper and lower hinges extend to the largest and smallest values, respectively but no further then 1.5XIQR from the hinge. P are Mann-Whitney U test. See also Supplementary Figure 1.

### CMV seropositive adults have expansion of Tfh1 cells, and reduced proportions of activated Tfh2 cells during infection

We have previously shown that in this cohort of CHMI participants, early activation of Tfh2 cells was associated with anti-parasitic antibody induction [17]. In contrast, activation of Tfh1 cells was not associated with antibody levels, but was associated with increased antibody secreting cell development which may impair germinal centres by acting as a nutrient sink during infection [18]. To assess if CMV seropositivity impacted Tfh cell differentiation either before or during CHMI, we analysed the same data set, and stratified by CMV serostatus. Tfh cells were defined as all CXCR5^+^ cells, subsets identified based on CXCR3 and CCR6 expression, and activation measured by expression of PD1, CD38 and ICOS. Non-Tfh effector CD4^+^ cells (CXCR5^-^ CD4^+^ T cells) were also analysed (Supplementary Fig. 2 A). Within the Tfh cell population, CMV seropositive individuals had a significantly higher proportion of Tfh1 cells, and a significantly decreased proportion of Tfh2 cells, before and during CHMI (Fig. 3 B). This expansion towards Th1-like cells was not seen in non-Tfh effectors, suggesting that the expansion of Tfh1 cells in CMV seropositive individuals was not due to a systemic inflammatory phenotype (Supplementary Fig. 2 C). Importantly, amongst activated Tfh cells (analysed as either PD1^+^, ICOS^+^ or CD38^+^ Tfh cells), there was a significantly increased proportion of Tfh1 cells (PD1^+^) and a reduced proportion of Tfh2 cells (ICOS^+^ and CD38^+^) at day 14/15 (Fig. 3 C). Within the CD4^+^ T cell population, there was no difference in the proportion of Tfh cells, nor any major differences in CD4^+^ effector (CXCR5-) Th1-, Th2- and Th17-like populations between CMV infected and uninfected individuals (Supplementary Fig. 2 B-C). Further, there was no difference in magnitude of activation in Tfh cells, nor non-Tfh effector cells during CHMI, aside from an increase in Th2 cell activation at day 14 post-inoculation (Fig. 3 A, Supplementary Fig. 2 D-E). While previous studies have shown that CMV antigen specific Tfh cells are dominated by Tfh1 cell subsets [36], these findings show that latent CMV infection modulates the entire Tfh cell compartment.

**Figure 3.**
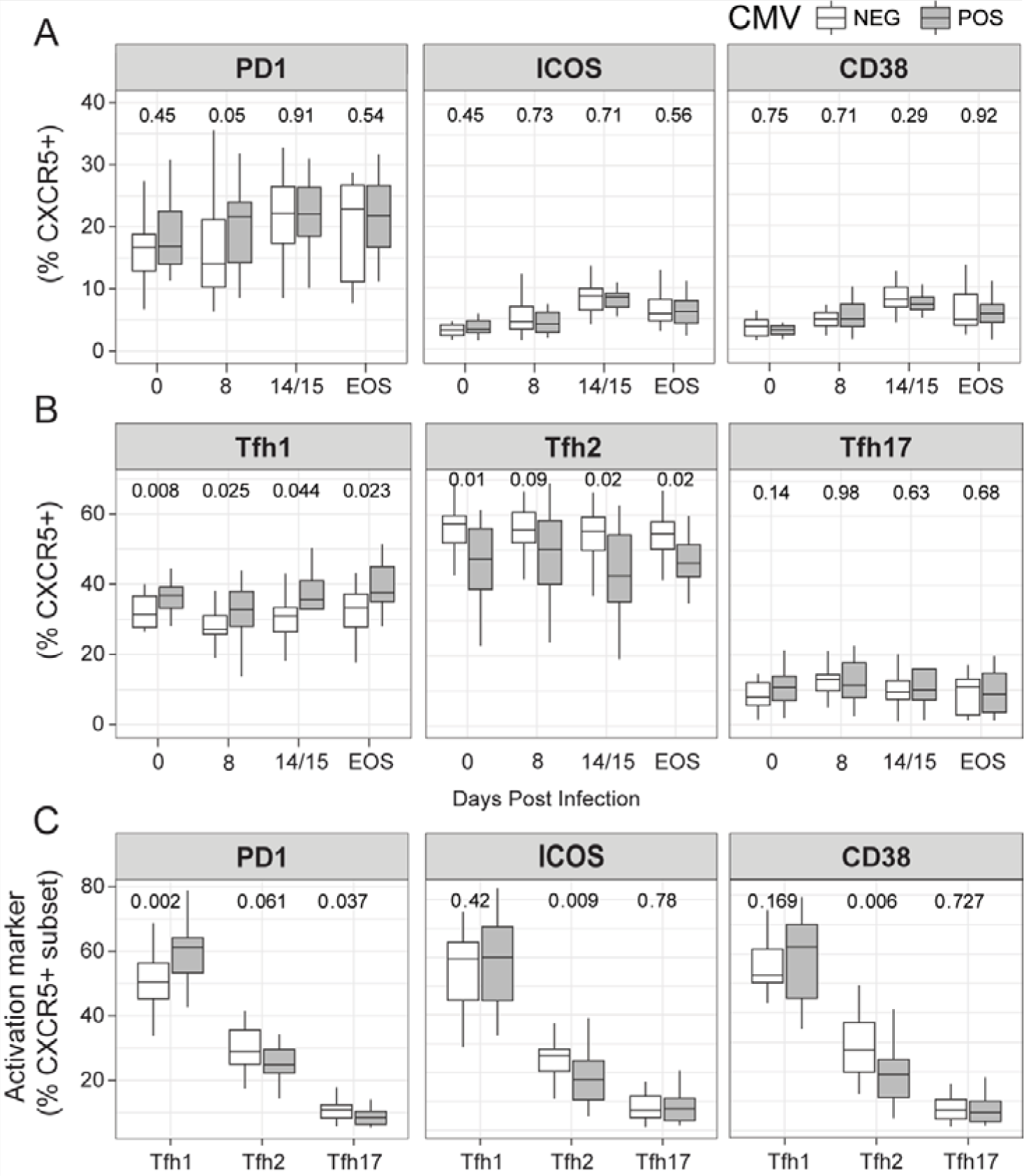
Tfh cell response is skewed to Tfh1 subsets in CMV seropositive adults. **(A)** Activation of Tfh cells as measured by PD1+, ICOS+, and CD38+ cells (% of Tfh CD4 T cells) in CHMI stratified by CMV status. **(B)** Tfh cell subsets (% of Tfh CXCR5+ CD4 T cells), stratified by CMV status. **(C)** Tfh cell subsets as a proportion of activated Tfh cells, stratified by CMV serostatus at day 14/15. CMV-white bars (n=19 for day 0, 8, 15, n=17 for day EOS), CMV+ grey bars (n=21 for day 0, 8, 15 and n=20 for EOS), data is Tukey boxplots with the median, 25^th^ and 75^th^ percentiles. The upper and lower hinges extend to the largest and smallest values, respectively but no further then 1.5XIQR from the hinge. P are Mann-Whitney U test. See also Supplementary Figure 2.

### The relationship between Tfh cell subsets and antibody induction is modulated by CMV serostatus

Different subsets of Tfh cells have specific functions within the germinal centre to drive B cell activation [15]. To assess if the CMV mediated bias of the Tfh cell compartment towards Tfh1 cells may impact antibody development in malaria, we assessed the correlations between Tfh cell subsets and antibody score in our cohort, stratified by CMV serostatus. The activation of Tfh2 cells, previously shown to be associated with antibody induction [17], was only associated with antibody score in CMV seronegative individuals (Fig. 4 A). Further, amongst CMV seropositive individuals, the proportion of Tfh1 cells in the Tfh cell compartment negatively correlated with antibody score (Fig. 4 B). Together, these data show that latent CMV infection modulates the structure of the Tfh cell compartment, which negatively impacts antibody production during malaria.

**Figure 4.**
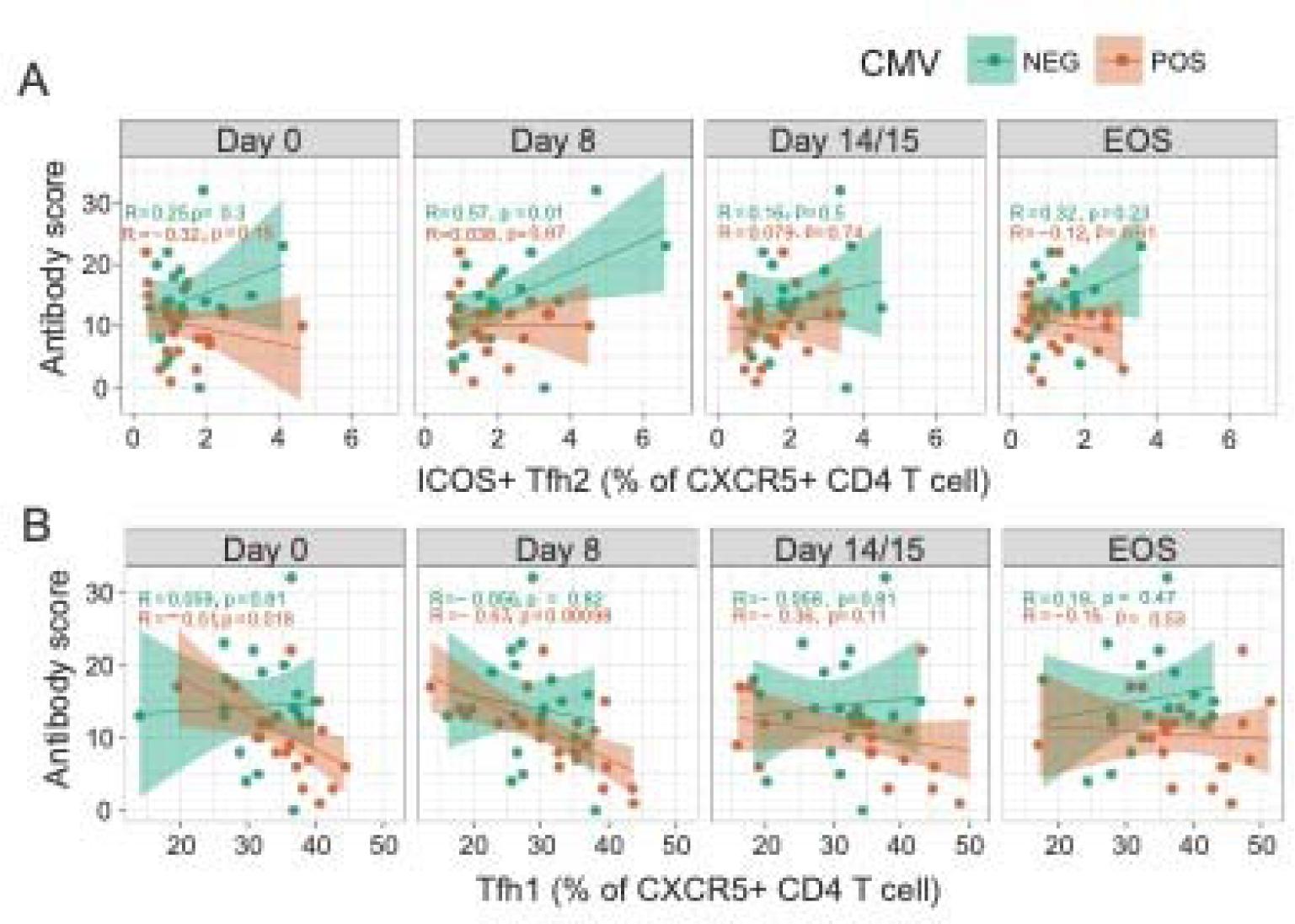
Correlation between Tfh cell subsets and antibody development based on CMV serostatus. **(A)** Correlation between activated Tfh2 cells (ICOS+ Tfh2 (% of Tfh CXCR5+ CD4T cells) and **(B)** the proportion of Tfh1 (% of Tfh CXCR5+ CD4 T cells) at day 0, 8, 14/15, EOS and Antibody score measured at EOS. CMV seronegative green (n=19 for day 0, 8, 15, n=17 for day EOS), CMV seropositive orange (n=21 for day 0, 8, 15 and n=20 for EOS). Spearman’s Rho and p are indicated.

### Latent CMV infection reduces antibody response to malaria vaccination

To assess if latent CMV infection also affects malaria vaccine responses, we analysed antibody levels in 24 individuals vaccinated with full-length MSP1 formulated with GLA-SE in a Phase 1a clinical trial [30,31]. Within this study 12 individuals were CMV seronegative and 12 were CMV seropositive. Sex, age and EBV status were comparable between CMV seropositive and seronegative individuals (Supplementary Table 1). We analysed induced MSP1 antibodies at day 85 following vaccination, which has been shown to be when antibody response peak [30,31]. IgG and IgM to MSP1, C1q fixation, opsonic phagocytosis by THP1 and neutrophils, and neutrophil antibody dependent respiratory burst (ADRB) responses were assessed. Additionally, IgG subclasses and NK IFNγ production and degranulation (measured by CD107α expression), were available for 11-13 participants. Unbiased clustering of all antibody responses aggregated the cohort into two clusters, low responders who were 8/12 (66%) CMV seropositive, and high responders who were 4/12 (33%) CMV seropositive, consistent with a negative impact of latent CMV infection on vaccine induced antibodies (Fig. 5A). To evaluate whether there was a difference in the antibody response between CMV positive and negative individuals, we used a MANOVA test which includes all antibody data in a single analysis. This found that there was a significant difference between CMV serostatus groups based on antibody parameters (MANOVA p=0.027, Fig. 5B). MSP1 targeted IgG, IgM, C1q and neutrophil ADRB were all reduced in CMV seropositive individuals. Differences were most pronounced for C1q fixation (Wilcox p=0.045, Fig. 5B). For antibody responses where only partial data was available, there was reduced IgG1, IgG3, IgG4, along with reduced NK IFNγ and degranulation in CMV seropositive individuals (Fig. 5C).

**Figure 5.**
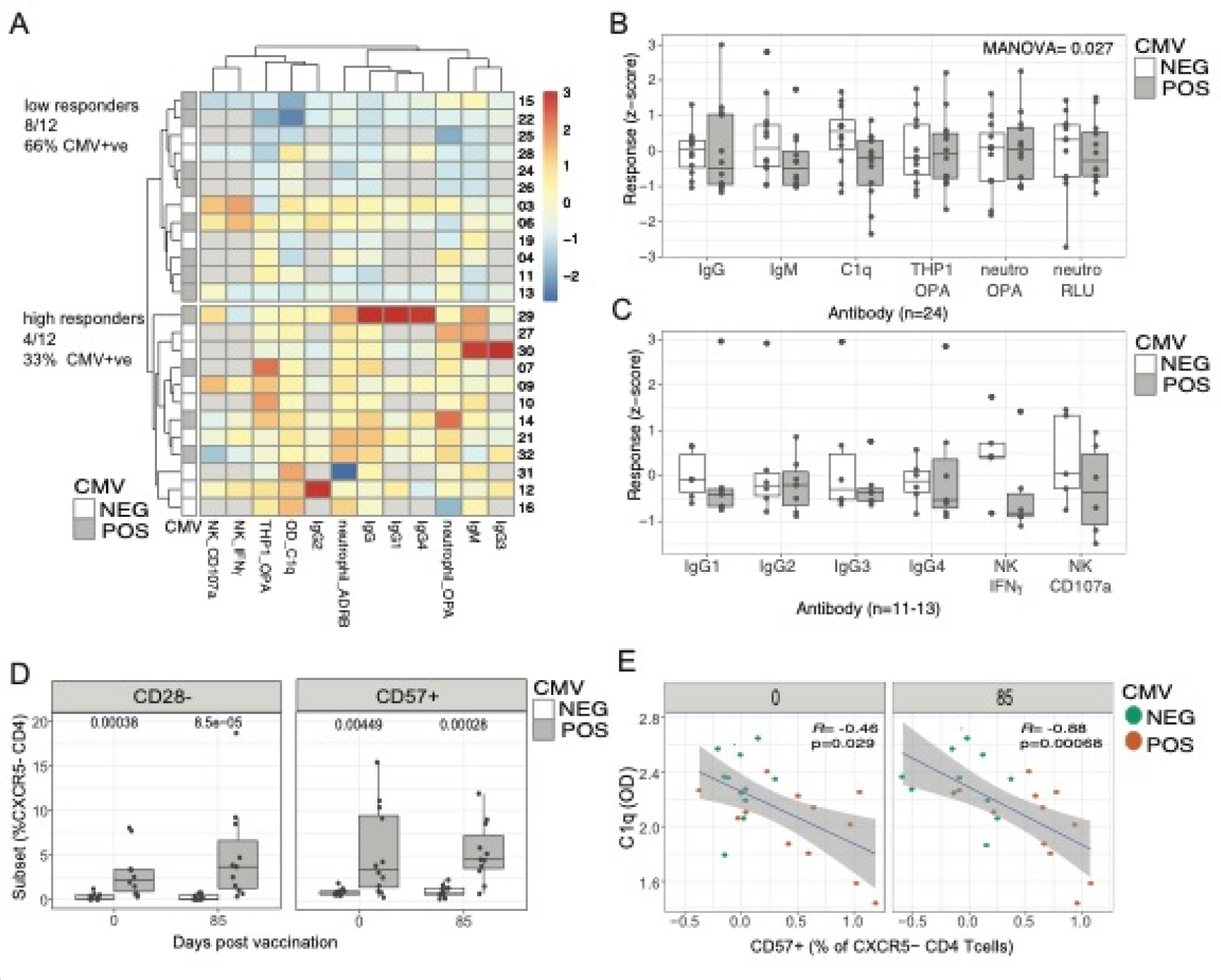
Impact of CMV serostatus on antibody responses to MSP1 vaccination. **(A)** Clustering of individuals within vaccination cohort based on MSP1 vaccine inducted antibodies. Antibody responses were normalised and z-scores to IgG, IgM and IgG1, 2, 3, and 4 subclasses, along with C1q complement fixation, opsonic phagocytosis (OPA) from THP1 cells and neutrophils, neutrophil antibody dependent respiratory burst (ADRB), and NK production of IFNγ and degranulation (CD107α). Individuals aggregated into low and high responders. CMV status is shown (n=24). **(B)** Impact of CMV on antibody responses for IgG, IgM, C1q, THP1 OPA, neutrophil OPA and neutrophil ADRB (n=24). Manova p is shown. (**C)** IgG subclasses, and NK responses between CMV negative and positive individuals (n=11-13). (**D)** Proportion of CD28- and CD57+ non-Tfh CD4 T cells (CXCR5-) at day 0 and day 85 stratified by CMV infection status. (**E)** Correlation between frequency of CD57+ non-Tfh CD4 T cells at day 0 and day 85, and C1q antibodies at day 85 following vaccination. Spearman’s Rho and p are indicated. See also Supplementary Figure 3.

To assess if CMV mediated cellular changes contributed to the reduced MSP1 antibody induction, we analysed the cellular compartment in vaccinated individuals at time of vaccination (day 0) and peak antibody response (day 85) (Supplementary Figure 3A). Unlike in the CHMI cohort, we did not detect any significant differences in subset distribution based on CMV infection status within Tfh and non-Tfh CD4^+^ T cells (Supplementary Figure 3B). Previous studies have reported CMV to be associated with down regulation of co-stimulatory marker CD28, and upregulation of terminally differentiation marker CD57 [25], both of which were included in cellular analysis. Consistent with previous studies, CMV infected individuals had increased frequencies of CD28^-^ CD4^+^ T cells, and increased frequencies of CD57^+^ CD4^+^ T cells within our cohort, both at day 0 and day 85 (Fig. 5D). These differences were significant within CXCR5^-^ CD4^+^ T cells, but not within the Tfh cell compartment (Supplementary Figure 3C). The frequency of CD57^+^ CD4 T cells at both day 0 and day 85 was negatively associated with the levels of C1q fixation, consistent with a role of CMV mediated terminally differentiation in reduced antibody responses to vaccination. Taken together, the data shows that latent CMV infection can modulate both the Tfh and non-Tfh CD4^+^ T cell compartments and that these changes negatively impact antibody production during malaria or following vaccination.

## Discussion

Here, we show that CMV serostatus has a substantial negative impact on antibody development during malaria infection and following vaccination. Within our CHMI cohort, CMV seropositive individuals had a bias in Tfh cell compartment towards Tfh1 cells, with the proportion of Tfh1 cells being negatively associated with antibody development. In our vaccination cohort, CMV seropositive individuals had increased frequencies of CD28^-^ and CD57^+^ CD4^+^ T cells, which was negatively associated with antibody development. These findings have significant implications for immune development during malaria via infection or vaccination in endemic areas, where there is typically a very high CMV seroprevalence [28,32].

CHMI studies are a powerful platform to evaluate mechanisms of immunity to malaria. In previous studies using this model, we have identified that expression of specific miRNAs prior to CHMI can predict individual control of parasite growth and antibody development following drug treatment [40], suggesting that preexisting host factors have major roles in the immune response to malaria. Here, we show that the protective role of Tfh2 cells in antibody induction [17] is significantly modulated by CMV serostatus, thus identifying a host factor underpinning immune heterogeneity. The impact of CMV on antibody induction was linked to a bias towards Tfh1 cells within the Tfh cell compartment. This link between CMV and modified Tfh cell compartment may explain previous reports that indicate that the impact of CMV on immune development in response to vaccination and infection is highly pathogen and/or context dependent. For example, while the impact of CMV serostatus on antibody induction to influenza vaccination is well studied, the findings are highly variable. Some studies have reported a negative impact of CMV seropositivity [41–48], others no impact [44,49–51], and others reporting a positive impact of CMV on antibody induction [52–54]. The impact of CMV on immune induction in the context of infection is also mixed, with antibodies induced by H1N1 influenza infection higher in CMV positive individuals [55], but no CMV mediated differences in antibodies induced following SARS-CoV-2 infection [56]. The role of Tfh cells in antibody induction is pathogen and context dependent [15]. In malaria, where Tfh2 cells have been associated with antibody development in response to infection [17], including in children in endemic areas [20], or after vaccination with both the licenced RTS,S [37] and experimental blood stage malaria vaccines [38,39], latent CMV infection may inhibit antibody induction. In contrast, following influenza vaccination, where Tfh1 cells are consistently positively associated with antibody induction [57,58], CMV seropositive individual may have enhance antibody development, as seen in some studies [52–54].

Highlighting the important role of our findings with CHMI to vaccine development for malaria, we also show that antibodies induced by full-length MSP1 vaccination are reduced in CMV seropositive individuals within a Phase 1 a clinical study. Despite the limited cohort size within this study, significantly reduced induction of complement fixing antibodies was shown in CMV seropositive individuals. While within the vaccine cohort we did not detected differences in the phenotype of Tfh cells, instead reduced vaccine induced antibodies were associated with increased frequencies of CD57^+^ CD4^+^ T cells, a cell subset thought to be terminally differentiated. The absences of detectible CMV mediated differences in the Tfh compartment may be due to the small cohort size, technical differences in Tfh subset staining, or may reflect genetic, environmental and past infection distinct between the CHMI and vaccine cohorts. Nevertheless, our findings are consistent with a recent study of immune responses induced by Ebola vaccination in UK and Senegalese participants [29]. In that study latent CMV infection was also associated with the reduced induction of antibodies following vaccination, and similarly changes to CD28 and CD57 expression within the T cell compartment were reported. Differences in CMV seroprevalence between recipients of this Ebola vaccine in the UK (50% CMV positive) and Senegal (100% CMV positive) accounted for the previously reported reduced vaccine responsiveness in the Senegalese cohort, with CMV infected UK participants having comparable responses to Senegalese individuals [59]. These differences in vaccine responsiveness in UK compared to Senegalese participants mimic differences in responsiveness to malaria vaccines seen between cohorts in the US and malaria endemic countries [6–10]. As such, studies to investigate the role of latent CMV infection on malaria vaccine responsiveness are required, particularly in target populations were CMV infection occurs early in life.

### Concluding remarks

Taken together, these data identify that latent CMV infection is an important driver of heterogeneity in the host immune response to malaria, with a striking negative effect on the development of malaria-specific antibodies induced by infection or vaccination. During malaria infection, this reduced antibody induction was linked to the skewing of Tfh cells to the Tfh1 cell subsets, and away from Tfh2 cells, which have previously been linked to antibody induction in CHMI [17]. In vaccination, reduced antibodies were associated with increased frequencies of CD57^+^ CD4^+^ T cells, highlighting a potential role of CMV mediated immune senescence in reduced antibody induction. These findings have important implications for understanding heterogeneity in immune responses to malaria infection and vaccination, particularly as CMV is typically highly prevalent and acquired in early life in areas of malaria transmission [28,32]. Further studies are required to assess if latent CMV infection contributes to the slow acquisition of protective functional antibodies in children in malaria endemic areas. Knowledge of the major impact of CMV on malaria immune responses may open new avenues for malaria control, such as targeting CMV infection or the immune changes induced by CMV infection to improve immunity to malaria and thus protection.

## Materials and Methods

### Study populations

Written informed consent was obtained from all participants. Ethics approval for the use of human samples in the relevant studies was obtained from the Alfred Human Research and Ethics Committee for the Burnet Institute (#225/19), the Human Research and Ethics Committee of the QIMR-Berghofer Medical Research Institute (P1479, P3444 and P3445) and the Ethics Committee of the Medical Faculty of Heidelberg (AFmo-538/2016) and the relevant regulatory authority (Paul Ehrlich Institute, Langen, Germany)

Controlled human malaria infection (CHMI) studies were performed as previously described using the induced blood stage malaria model [60]. Malaria naïve individuals were inoculated by intravenous injection of 2800 *P. falciparum* infected red blood cells and monitored for parasite growth with qPCR [61]. Here, blood samples were collected at baseline (day 0), day of treatment (day 8) and at 14 or 15 days and at end of study (EOS), 27-36 days after inoculation (4 studies, across 6 independent infection cohorts). Clinical trials were registered at ClinicalTrials.gov NCT02867059 [62], NCT02783833[63], NCT02431637 [64], NCT02431650 [64]. Whole blood was stained and analysed immediately by flow cytometry and plasma was collected from lithium heparin collection tubes.

Full-length MSP1 vaccination was performed as previously described in a Phase 1a clinical trial of full-length MSP1 adjuvanted with GLA-SE [30,31] registered with EudraCT (No. 2016-002463-33. Volunteers received three vaccinations at intervals of 29±3 days at 25, 50 or 150μg. For the current study, antibody responses at day 85 following first vaccination, and cellular responses at day 0 and day 85 were considered. Antibody responses were measured from sera collected in S-Monovetter serum-gel, and PMBCs were collected from sodium-citrate tubes.

### CMV and EBV serostatus

For CHMI cohorts CMV and EBV seroprevalence was assessed using baseline plasma samples by commercially available ELISA kits (ab108724 and ab108730), according to manufacturer’s instructions. CMV ELISA kit has a reported sensitivity and specificity of 98% and 97.5% respectively. For the vaccination cohort, CMV seroprevalence was assessed using day 0 plasma samples by commercially available CMV IgG ELISA (EUROIMMUN EI 2570-9601 G). One individual fell into a CMV intermediate status in the ELISA. For this sample, an immunblot (MIKROGEN DIANOSTIK *recom*Line CMV IgG 5572) was performed which was positive. The EBV seroprevalence was established using the Anti-EBV CA_ELISA (IgG) Assay (EUROIMMUN EI 2791-9601 G).

### Induced antibody responses

For CHMI studies, antibody response to intact merozoites and merozoite surface antigen MSP2 were quantified as previously described [17]. To isolate merozoites, *P. falciparum* 3D7 parasites were maintained in continuous culture in RPMI-HEPES medium supplemented with hypoxanthine(370 mM), gentamicin (30 mg/ml), 25 mM sodium bicarbonate and 0.25% AlbuMAX II (GIBCO) or 5% heat-inactivated human sera in O+ RBCs from malaria-naive donors (Australian Red Cross blood bank). Cultures were incubated at 37°C in 1% O_2_, 5% CO_2_, 94% N_2_ and schizont stage parasites were purified by MACS separation (Miltenyl Biotec). To isolate merozoites, magnet purified schizonts were incubation with the protease inhibitor E64 (10 mg/ml), and following complete development, merozoites were isolated by membrane filtration (1.2 mm). 50 ml of *P. falciparum* 3D7 merozoites (2.5 × 10^5^ merozoites/ml) or 50 ml of 0.5 mg/ml MSP2 recombinant antigen [65] in PBS were coated to 96-well flat bottom MaxiSorb plates (Nunc) overnight at 4°C. Plates were blocked with 10% skim milk for merozoites, or 1% casein (Sigma-Aldrich) for MSP2 for 2 hours at 37°C. Plasma was diluted in 0.1% casein in PBS1/100 for IgG, 1/250 for IgG subclasses and IgM, 1/100 for C1q, 1/100 for FcγR targeting merozoites or 1/50 for FcγR targeting MSP2) and incubated for 2 hours at room temperature. For total antigen specific IgG detection, plates were incubated with goat polyclonal anti-human IgG HRP-conjugate (1/1000; Thermo Fisher Scientific) for 1 hour at room temperature. For detection of IgG subclasses and IgM, plates were incubated with a mouse anti-human IgG1 (clone HP6069), mouse anti-human IgG3 (HP6050) or mouse anti-human IgM (clone HP6083) at 1/1000 (Thermo Fisher Scientific) for 1 hour at room temperature. This was followed by detection with a goat polyclonal anti-mouse IgG HRP-conjugate (1/1000; Millipore). For all ELISAs, plates were washed three times with PBS (for merozoite ELISAs) or PBS-Tween 0.05% (for MSP2) between antibody incubation steps. For detection of complement fixing antibodies, following incubation with human sera, plates were incubated with purified C1q (10 mg/ml; Millipore) as a complement source, for 30 min at room temperature. C1q fixation was detected with rabbit anti-C1q antibodies (1/2000; in-house) and a goat anti-rabbit-HRP (1/2500; Millipore). For FcγR assays, 100ul of biotin-conjugated rsFcγRIIa H131 or rsFcγRIIIa V158 ectodomain dimer (0.2ug/ml) was incubated at 37°C for 1 hour followed by 3 washes with PBS-Tween. The binding was detected with horseradish peroxidase (HRP)-conjugated streptavidin antibody (1:10,000) in PBS-BSA at 37°C for 1 hour.

TMB liquid substrate (Life Technologies) was added for 1 hour at room temperature and the reaction was stopped using 1M sulfuric acid. The optical density (OD) was read at 450 nm. For opsonic phagocytosis, THP1 cells were incubated with intact merozites (stained with Ethidium Bromide and opsonised with plasma diluted 1/100) or latex beads coated with MSP2 (opsonised with plasma diluated 1/10) for 20 min at 37°C and cells washed with FACS buffer. The proportion of THP-1 cells containing fluorescent-positive beads was evaluated by flow cytometry (FACS CantoII, BD Biosciences), analyzed using FlowJo software and presented as phagocytosis index (the percentage of THP-1 monocytes with ingested merozoites or beads).

To calculate antibody score antibody responses below positive cut-off threshold were set as negative, and remaining positive responses were used to calculate median and used to categorise responses into low (below median) and high (above median) responses. Antibody score was calculated by giving categories zero/low/high a numerical score of 0/1/2 and then summing across all antibody responses.

For full-length MSP1 vaccination studies, antibodies were measured as previously reported [30,31]. Total IgG and IgM antibody levels were determined by ELISA using MaxiSorp plates (Thermo Fisher Scientific) coated with 100nM recombinant full-length MSP1 (Glycotope Biotechnology GmbH, Heidelberg). Sera were titrated in two-fold dilutions and incubated for 2h. A secondary antibody goat anti-human IgG and IgM alkaline phosphatase conjugate (Sigma-Aldrich) was used at a dilution of 1:20,000 for 1h. The substrate p-nitrophenyl-phosphate was added and incubated for 1h in the dark, and stopped with 1M NaOH. The absorbance at 405 nm was determined using the plate reader Cytation 3 (BioTek). For the IgG subtype ELISA, the same protocol was applied, with the exception that subclass-specific peroxidase-conjugated secondary antibodies were used (The Binding Site GmbH). The substrate 1-step turbo TMB (Thermo Fisher) was added for 20 min in the dark, then stopped by adding 1 M HCl. Optical density was recorded at 450nm. For the C1q fixation ELISA assay, full-length MSP1 coated plates were also incubated overnight at 4°C. The next day, the plates were washed four times with 1x PBS containing 0.05% Tween 20 and blocked with 1% casein/PBS at 37°C for 2 h. After washing, the plates were incubated with 50µl of purified IgG at 1mg/ml blocking buffer, respectively at 37°C for 1 h. After incubation, the plates were washed and recombinant C1q (Abcam) at 10µg/ml in blocking buffer was added for 30 min at 37°C. To detect C1q binding, anti-C1q horse radish peroxidase (HRP)-conjugated secondary antibodies (Abcam) was added at a 1:100 dilution in blocking buffer for 1h at 37°C. Then SigmaFAST OPD (Sigma-Aldrich) was added for 30 min to 1h in the dark at room temperature for development. The reaction was stopped with 1M HCL and the signal intensity was measured at 492nm using the Biotek Cytation 3 plate. For the Antibody-dependent respiratory burst (ADRB) assay full length MSP1 protein was coated at 100nM in PBS in opaque 96-well Lumitrac microplates (Greiner Bio-One), blocked with 1% casein/PBS and incubated for 1h at 37°C with 50µ/well of IgG at 1mg/ml. The plates were washed with PBS and 50µl/well of luminol at 0.04mg/ml were added to each well. Next, freshly isolated neutrophils were added at 10^7^ cells/ml and absorption was immediately read at 450nm using the Biotek Cytation 3 reader for every 2 min over 1 h. For the Opsonic phagocytosis activity assay, full-length MSP1 coupled to fluorescent beads were performed as previously reported [30,31] using THP1 cells and neutrophils). Briefly, an antigen coupled bead suspension was added to 96-well to U-bottomed plates and opsonized for 1h at 37°C with 50µl/well of purified IgG. The plates were centrifuged at 2000x g for 7 min and washed with PBS. Opsonized beads were resuspended in culture medium (RPMI 1640 media with 2mM L-glutamine, 10% FCS and 1% penicillin-streptomycin) before incubation with 5 x 10^4^ phagocytes at 37°C. Phagocytosis was arrested by centrifugation at 1200 rpm for 7 min at 4°C and washing with ice-cold FACS buffer (0.5% BSA and 2mM EDTA in PBS). Cells were resuspended in 2% formaldehyde/PBS and the proportion of phagocytes containing PE fluorescent beads was on a FACS Canto II (BD biosciences). NK activity assays, full-length MSP1-coated plates were washed PBS and blocked with 1% casein/PBS for 4 h at 37°C. The plates were washed and incubated for 2h at 37°C with IgG at 1mg/ml. Then 5 x 10^4^ freshly isolated human NK cells together with anti-human CD107a PE (BD biosciences,1:70), brefeldin A (Sigma-Aldrich, 1:200) and monensin (Sigma, 1:200) was added for 18 h at 37°C. Afterwards, the cells were transferred to 96-well V-bottomed plates, centrifuged at 1500 rpm for min at 4°C and washed with ice-cold FACS buffer. Cell viability was determined by adding dye eFluor^TM^520 (Thermo Fischer). NK cell surface markers were labelled with an antibody cocktail of anti-CD56 APC (BD biosciences, 1:17) and anti-CD3 PE-Cy5 (BD biosciences, 1:33) for 30 min at 4°C in the dark. After washing, NK cells were fixed with CellFIX (BD biosciences) at 4°C and permeabilized with permwash (BD biosciences) for 10 min at 4°C. Intracellular IFN-γ was measured by adding anti-IFN-γ PE-Cy7 (BD biosciences, 1:33) for 1h at 4°C. NK cells were washed with permwash, resuspended in FACS buffer and Ab-NK activity (proportion of NK cells with CD107a and/or IFN-γ staining was assessed by a FACS ContoII (BD biosciences).

### Quantification of T cells

For CHMI studies, whole blood was analysed as described previously [17]. 200 mL of whole blood were stained with the following conjugated antibodies, all from BD Biosciences; anti-CD20-BUV395 (2H7, 1:150, Cat# 563782), anti-CXCR5-BV421 (RF8B2, 1:50, Cat#562747), anti-CD4 (V500, 1:30, Cat# 560768), anti-CCR6-BV650 (11A9, 1:200, Cat#563922), anti-CD38-BV786 (HIT2, 1:400, Cat#563964), anti-CXCR3-APC (1C6, 1:25, Cat#550967), anti-CD27-APC-R700 (M-T271, 1:100, Cat#565116), anti-CD8-APC-Cy7 (SK1, 1:150, Cat#557834), anti-CD19-FITC (HIB19, 1:20, Cat#55412), anti-CD45-PerCP-Cy5.5 (2D1, 1:50, Cat#340953), anti-ICOS-PE (DX29, 1:10, Cat#557802), anti-CD3-PE-CF594 (UCHT1, 1:600, Cat#562280), anti-PD1-PE-Cy7 (EH12.1, 1:100, Cat#561272). RBCs were lysed with FACS lysing solution (BD) and re-suspended in 2% FBS/PBS. Samples were acquired on the BD LSR Fortessa TM 5 laser cytometer (BD Biosciences). These data were analyzed using FlowJo version 10.8 software (Tree Star, San Carlos, CA, USA).

For full-length MSP1 vaccine cohorts, cellular responses were analysed as previously reported [31]. The phenotyping of T- and B-lymphocytes was conducted in whole blood samples within 24 hours. To monitor the dynamics of lymphocyte subpopulations, comprehensive T-cell and B-cell panels were employed, based on modified recommendations from the “Human Immunophenotyping Consortium” [66].

## Data availability statement

The data underlying all figures are available in the published article and its online supplemental material.

## Funding and financial conflicts of interest

Authors declare no conflicts of interest.

RTL is employee from Sumaya GmbH & Co. KG.

## Acknowledgements

RBC and human serum were provided by the Australian Red Cross Blood Bank (Melbourne). We thank the participants involved in the controlled human malaria infection studies and all study clinicians and support staff at QPharm and Medicine for Malaria Venture for funding the CHMI studies. This work was supported by the National Health and Medical Research Council of Australia (program grant 1132975 to J.S.M. and C.R.E.; Practitioner Fellowship 1135955 to J.S.M., Senior Research Fellowship 1154265 to C.R.E., Career Development Award 1141278, Project Grant 1125656, and Ideas Grant 1181932 to M.J.B.; Program Grant 290208, Senior Research Fellowship 1077636 to J.G.B.; Emerging Leadership 2 Fellowship to BEB, Australian Centre for Research Excellence in Malaria Elimination 1134989 to J.G.B. and J.S.M.); the Jim and Margaret Beever Fellowship to J.A.C; the CSL Centenary Fellowship to M.J.B, and the Snow Medical Foundation Fellowship to M.J.B. The Burnet Institute is supported by the NHMRC for Independent Research Institutes Infrastructure Support Scheme and the Victorian State Government Operational Infrastructure Support.

Special thanks to the volunteers that participated in the first-in-human phase 1a clinical trial with full-length MSP1 in Heidelberg and to all the team at the Parasitology Unit as well as the Clinical Pharmacology and Pharmacoepidemiology Unit at the University Hospital Heidelberg, for their enthusiastic work in running the trial and ulterior analysis of samples. This work was funded by Sumaya Biotech GmbH & Co. KG, Heidelberg, Germany, via a venture loan provided by the EU Malaria Fund. We are very thankful to the entire team at Sumaya-Biotech for constant support.

