## Supplementary Tables and Figures for "Heterogeneity of the human immune response to malaria infection and vaccination driven by latent cytomegalovirus infection"

**Supplementary Table 1: Vaccine cohort demographics**

|  | CMV serostatus | |
| --- | --- | --- |
|  | Negative | Positive |
| Total, n (%) | 12 (50%) | 12 (50%) |
| Sex, male, n (%) | 6 (50%) | 4 (33.3%) |
| EBV, positive, n (%) | 8 (66.6%) | 12 (100%) |
| Age years, median [IQR] | 25, [22-42.5] | 28, [23.75-38.25] |

**Supplementary Figures**

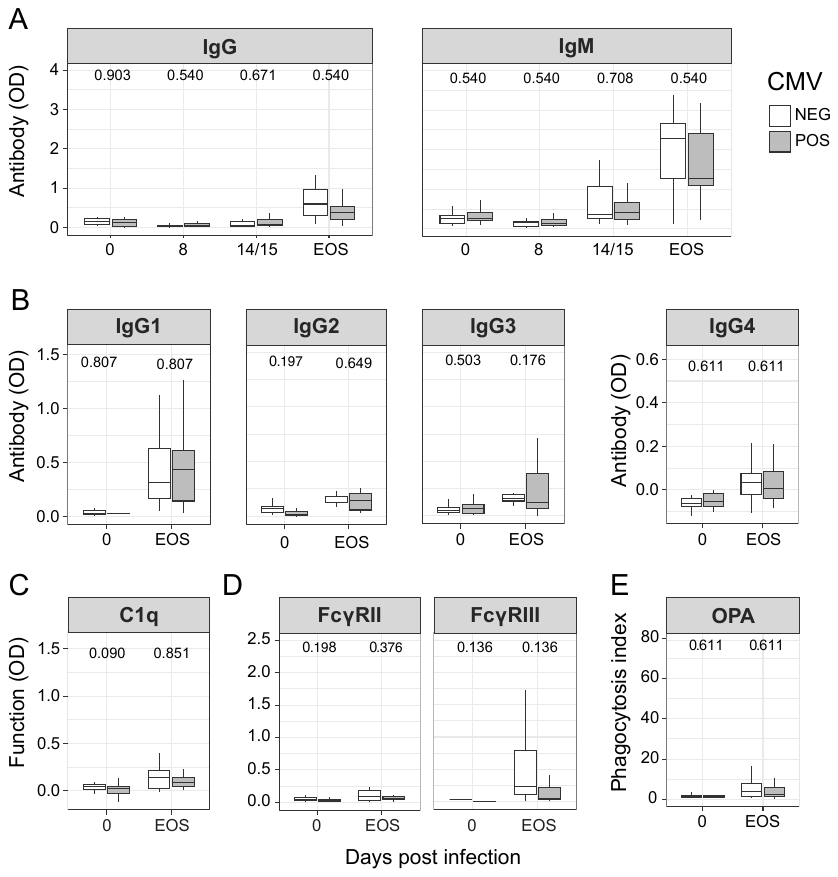

Supplementary Figure 1. **Antibodies to merozoite surface protein 2 induced by controlled human malaria infection stratified by CMV infection status**

**(A)** IgG and IgM antibodies, **(B)** IgG subclass antibodies and levels of functional antibodies that can **(C)** fix C1q complement component, **(D)** cross link FcgRII and FcgRIII, and **(E)** drive opsonic phagocytosis by THP1 cells, targeting merozoite surface protein 2 antigen, stratified by CMV serostatus. CMV-ve white bars (n=19), CMV+ grey bars (n=21), data is Tukey boxplots with the median, 25^th^ and 75^th^ percentiles. The upper and lower hinges extend to the largest and smallest values, respectively but no further then 1.5XIQR from the hinge. P are Mann-Whitney U test.

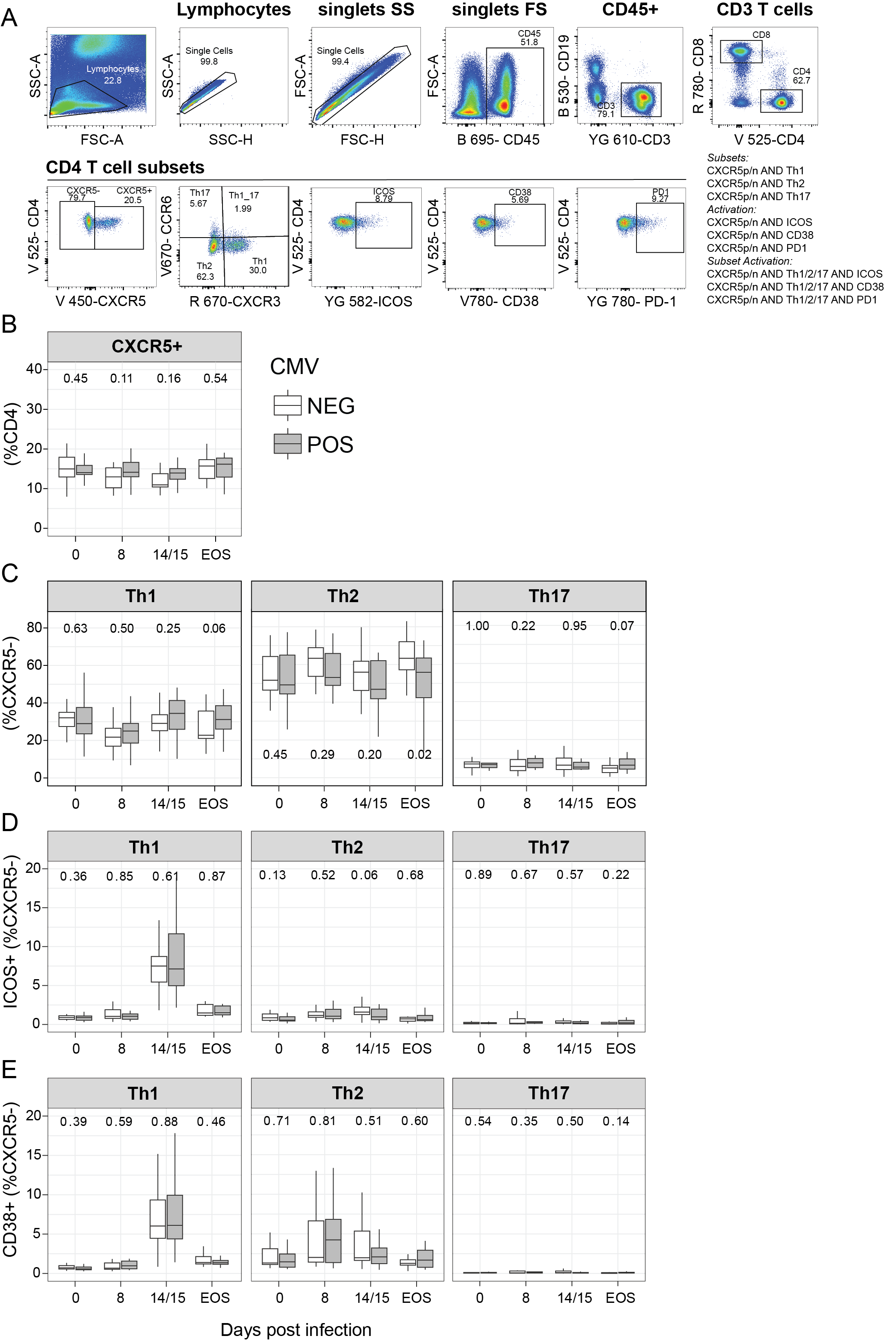
Supplementary Figure 2: **Limited impact of CMV serostatus on non-Tfh effector cell subsets**

**(A)** Gating strategy to identify Tfh cells, subsets and activation status. Tfh cells were identified as CXCR5+ cells. Subsets were identified as Tfh1 – CXCR3+CCR6-, Tfh2 – CXCR3-CCR6-, Tfh17 – CXCR3-CCR6+. Activation status was gated as PD1+, ICOS+, CD38+. Activation on Tfh and non-Tfh cells (CXCR5-) was analysed with AND gating.

**(B)** Proportion of Tfh cells as a percent of CD4 T cells in CHMI stratified by CMV status. **(C)** T helper cell subsets (CXCR5-) as a proportion CD4 T cells, stratified by CMV serostatus. **(D)** Proportion of ICOS+ activated T helper cell subsets (CXCR5-) and **(E)** CD38+ activated T helper cell subsets (CXCR5-), stratified by CMV serostatus. CMV-ve white bars (n=19 for day 0, 8, 15, n=17 for day EOS), CMV+ grey bars (n=21 for day 0, 8, 15 and n=20 for EOS), data is Tukey boxplots with the median, 25^th^ and 75^th^ percentiles. The upper and lower hinges extend to the largest and smallest values, respectively but no further then 1.5XIQR from the hinge. P are Mann-Whitney U test.

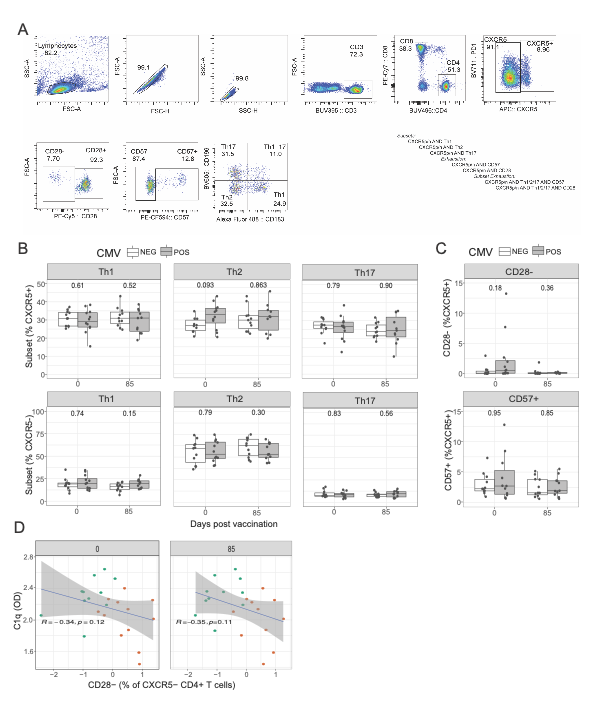

Supplementary Figure 3: **Analysis of cellular compartment in MSP1 vaccinated individuals.**

**(A)** Gating strategy to identify helper subsets within Tfh (CXCR5+) and non-Tfh (CXCR5-) CD4+ T cells. CD4+ T cells were identified as CXCR5+ (Tfh) or CXCR5- (non-Tfh) cells. Subsets were identified as Th1 – CXCR3+CCR6-, Th2 – CXCR3-CCR6-, Th17 – CXCR3-CCR6+. Potential exhaustion status was analysed based on CD28 and CD57 expression. Subsets and exhaustion on Tfh and non-Tfh cells was analysed with AND gating. **(B)** T helper subset distribution in CXCR5+ and CXCR5- CD4 T cells at day 0 and day 85. **C)** CD28- and CD57+ frequencies in CXCR5+ Tfh cells at day 0 and day 85. For B and C Wilcox P value is indicated. **D)** Correlation between CD28+ CXCR5- CD4 T cells at day 0 and day 85 and C1q antibodies induced at day 85. Spearmans Rho and p are indicated.
